# Fluctuating chromatin facilitates enhancer-promoter communication by regulating transcriptional clustering dynamics

**DOI:** 10.1101/2024.08.30.610578

**Authors:** Tao Zhu, Chunhe Li, Xiakun Chu

## Abstract

Enhancers regulate gene expression by forming contacts with distant promoters. Phase-separated condensates or clusters formed by transcription factors (TFs) and co-factors are thought to facilitate these enhancer-promoter (E-P) interactions. Using polymer physics, we developed distinct coarse-grained chromatin models that produce similar ensemble-averaged Hi-C maps but with “stable” and “dynamic” characteristics. Our findings, consistent with recent experiments, reveal a multi-step E-P communication process. The dynamic model facilitates E-P proximity by enhancing TF clustering and subsequently promotes direct E-P interactions by destabilizing the TF clusters through chain flexibility. Our study promotes physical understanding of the molecular mechanisms governing E-P communication in transcriptional regulation.

**Graphical TOC Entry:** 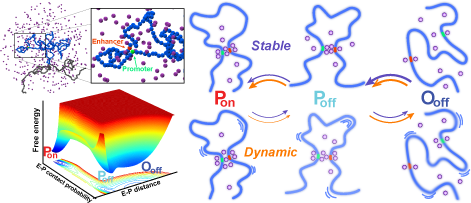

---

Precise spatio-temporal regulation of gene expression patterns is governed by DNA regulatory elements, particularly enhancers.^1–4^ Enhancers can initiate transcription by activating target gene promoters, which are often located thousands to millions of base pairs away. ^5,6^ It is well established that these distantly-separated enhancers and promoters in the 1D genomic sequence must come into close proximity in 3D nuclear space for gene expression.^7^ However, understanding the underlying mechanisms of how the long-range enhancer-promoter (E-P) interactions are efficiently established *in vivo* remains a central challenge in genome biology. ^8^

Significant efforts have been made to characterize the long-range E-P interactions that controls transcriptional activities.^9–12^ Over the past decades, techniques such as chromosome conformation capture (3C) and its derivatives, including Hi-C,^13^ have revealed the hierarchical organization of the 3D genome, showing well-organized E-P contacts.^14^ At the megabase scale, the genome is folded into self-interacting regions known as topologically associating domains (TADs), whose boundaries are determined by the binding of CTCF or other insulator proteins.^15–17^ Enhancers and their target promoters frequently reside within the same TAD, providing a confined space for DNA segments to interact, thus facilitating the establishment of E-P contacts.^3,18^ While numerous experiments have shown that disruptions in TAD organization can break E-P contacts,^19–22^ others have observed that E-P contacts can be independent of TAD boundaries and alterations in TAD structures do not significantly affect the expression of most genes.^23–26^ This complex interplay between TAD organization and E-P contact highlights the need for further investigation into the physical mechanisms underlying E-P communication.

Recent Hi-C experiments have shown that TAD boundaries are highly conserved during cell differentiation, even as gene expression patterns change significantly.^27^ Meanwhile, extensive Hi-C data analysis has revealed that E-P contacts can persist across different cell types, regardless of whether the target genes are expressed,^28,29^ leading to a complex relationship between E-P contacts and gene expression. An emerging model that may explain the uncoupling of E-P contacts from transcriptional activities involves transcriptional clusters or phase-separated condensates induced by transcription factors (TFs) and other co-activators.^30–33^ These membraneless organelles are observed throughout the nucleus and create microenvironments with high local concentrations of TFs, facilitating E-P spatial proximity.^34,35^ Subsequently, non-random specific E-P interactions are established to control gene transcription.^5^

The efficiency of this “facilitating-specifying” model may be modulated by the stability-dynamics balance of TF condensates. ^36,37^ While these clusters promote the initial step by reducing the search space for E-P communication, they may hinder subsequent E-P contact formation due to the highly crowded molecular assembly. On the other hand, since Hi-C data reflects the ensemble-average properties of chromatin contacts, similar E-P contact maps can arise from different chromatin structural fluctuations.^38^ It is acknowledged that chromatin dynamics and structural heterogeneity play crucial roles in precisely regulating gene expression, considering cell-to-cell variability.^39^ However, how the fluctuations of chromatin domains, in conjunction with TF clustering, regulate E-P contacts during cell-state transitions remains unclear.

In this Letter, we developed two coarse-grained (CG) molecular dynamics (MD) simulation chromatin models with varying degrees of structural fluctuations to study E-P communication involving TF clustering. The CG models are based on a beads-on-a-string representation, with each bead representing a 30-nm chromatin fiber corresponding to a 3-kbp DNA fragment.^40,41^ We focused on a TAD with a length of 1 Mbp, which represents the typical average length of TADs (Figure 1A).^15^ Within the TAD, the system includes a pair of enhancer and promoter, separated by a genomic distance of 120 kbp, close to the experimentally measured averages.^42^ The effect of cohesin protein was implemented as a harmonic potential working at the boundary points of the TAD. These two models, which exhibited distinct chain dynamics, were constructed using pairwise interactions (*f_i,j_*) derived directly from the Lennard-Jones (LJ) potential and determined by the maximum entropy principle (MEP) iteration.^43–45^ We refer to these as the “stable” and “dynamic” models, respectively (Figure 1A-C; details in Supporting Information).

**Figure 1:**
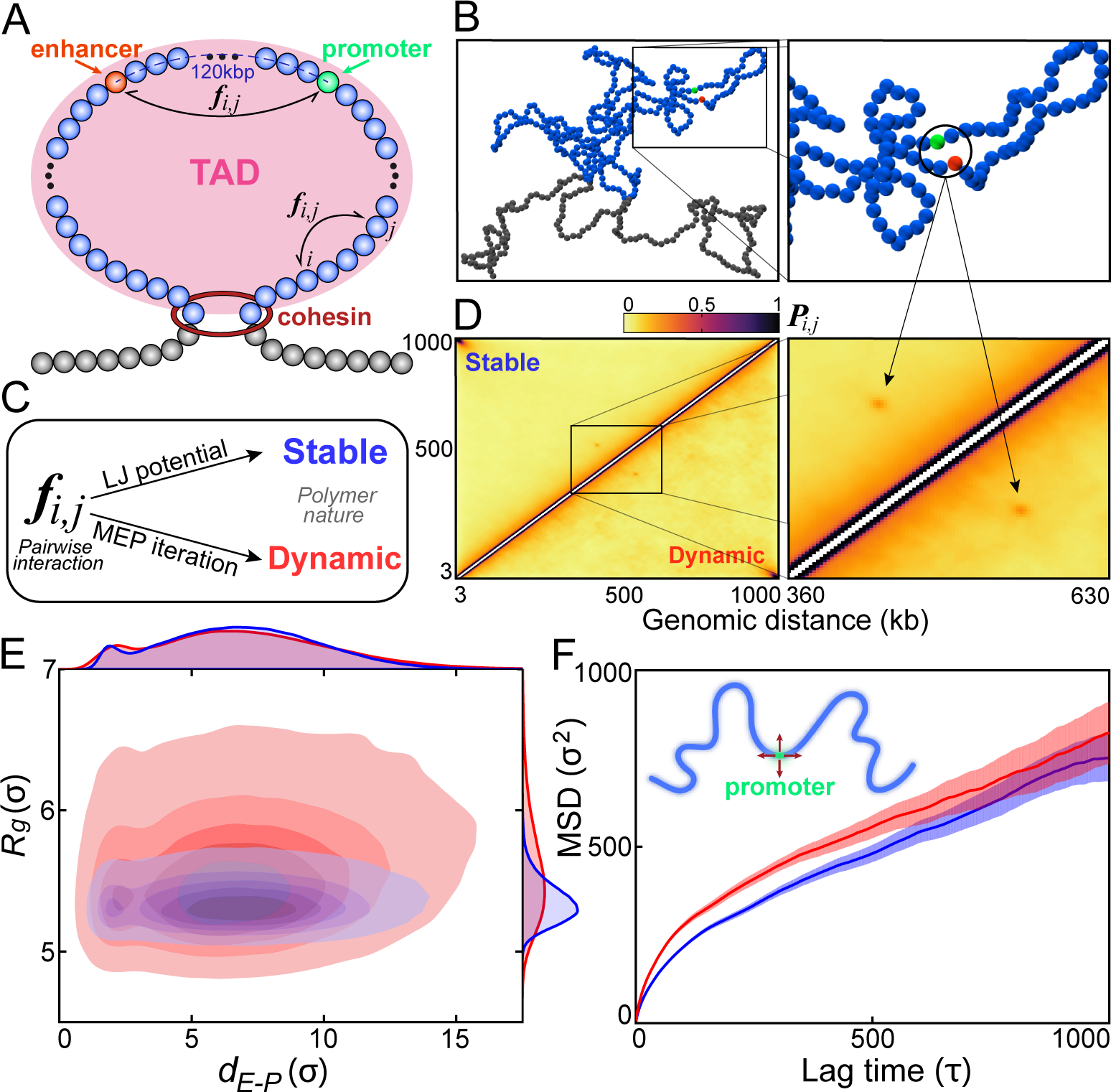
Structural and dynamic properties of stable and dynamic chromatin models. (A) Schematic illustration of the coarse-grained (CG) chromatin model, representing a TAD with a length of 1 Mb. Chromatin within the TAD is colored blue, while chromatin outside the TAD is colored gray. The enhancer and promoter are each represented by a single bead, colored red and green, respectively. Cohesin protein, illustrated as a red ring, is modeled as a distance-restraint potential acting on the boundary points of the TAD. (B) A simulation snapshot showing the enhancer and promoter in close contact. (C) Stable (blue) and dynamic (red) chromatin models with pairwise interactions (*f_i,j_*) generated by directly using the Lennard-Jones (LJ) potential and through the maximum entropy principle (MEP) iteration, respectively. (D) Hi-C-like contact maps (*P_i,j_*, contact probability formed by loci *i* and *j*) for the stable and dynamic chromatin models, showing high similarity. (E) Distributions of the radius of gyration (*R_g_*) of the chromatin system and spatial distance between the enhancer and promoter (*d_E−P_*) for the stable and dynamic models. (F) Mean squared displacement (MSD) of the promoter as a function of lag time for the stable and dynamic chromatin models. *σ* is the reduced unit of length, representing the diameter of the CG bead and *τ* is the reduced unit of time (details in Supporting Information).

Despite the high similarity between the ensemble-average Hi-C-like contact maps gen-erated by these two models (Figure 1D and Figure S1), the structural distributions of the chromatin ensembles differ significantly (Figure 1E). The result showed that the dynamic model explores a broader conformational space than the stable model in terms of overall TAD compaction degree (*R_g_*, radius of gyration) and the E-P distance (*d_E−P_*). To further characterize the chain fluctuations in these two chromatin models, we calculated the mean squared displacement (MSD) of the promoter region as a function of lag time (*t*): 〈(**r***_c_* (*t*_0_ + *t*) *−* **r***_c_* (*t*_0_))^2^〉*_t_*_0_, where **r***_c_* (*t*_0_) is the position of the center of mass of the region at simulation time *t*_0_ (Figure 1F). Although both models exhibited sub-diffusive behavior for the beads within the chromatin domain (Figure S1), consistent with previous simulation and experimental observations,^46–51^ the promoter in the dynamic model diffuses faster than in the stable model, indicating a more fluctuating chain dynamics. Consequently, the structural ensembles of chromatin generated by the dynamic model are more heterogeneous and associated with higher chain fluctuations compared to the stable domain, even though both models exhibited similar structural behavior from the Hi-C perspective.

TFs can specifically bind to sequences of enhancers and promoters, promoting DNA folding and the formation of E-P loops.^52,53^ This TF binding brings distal enhancers into physical proximity with target gene promoters, facilitating direct E-P interactions. The resulting 3D chromatin structures are critical for gene regulation, and alterations in these structures, involving TF-mediated E-P communication, can drive cell-fate decisions.^54^ To study the impact of chromatin dynamics on TF-mediated E-P communication that subsequently impacts gene expression, TFs were explicitly incorporated into the chromatin chain system represented by stable and dynamic models (Figure 2A and 2B). To accurately capture the effects of chromatin dynamics on E-P interactions and the role of TF clustering in this process, TFs were modeled as CG beads attractively interacting with both enhancers and promoters through LJ potential, but with only volume-excluding interactions among them.

We observed an increase in the E-P contact probability (*p_E−P_*) with the addition of TFs into the system (number of TFs, *n_TF_*) for both chromatin models (Figure 2C), highlighting the role of TFs in promoting E-P contact formation. However, *p_E−P_* exhibited distinct behaviors with respect to the changes in *n_TF_* for these two models. For smaller values of *n_TF_* (i.e., *n_TF_ <* 200), the relationship between *p_E−P_* and *n_TF_* is similar for both models. While with a larger number of TFs, the dynamic model showed a more significant increase in *p_E−P_* with increasing *n_TF_* compared to the stable model. To better understand the relationship between *p_E−P_* and *n_TF_*, we fitted the data using the Hill function (details in Supporting Information):^55^

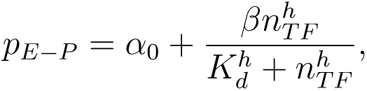

**Figure 2:**
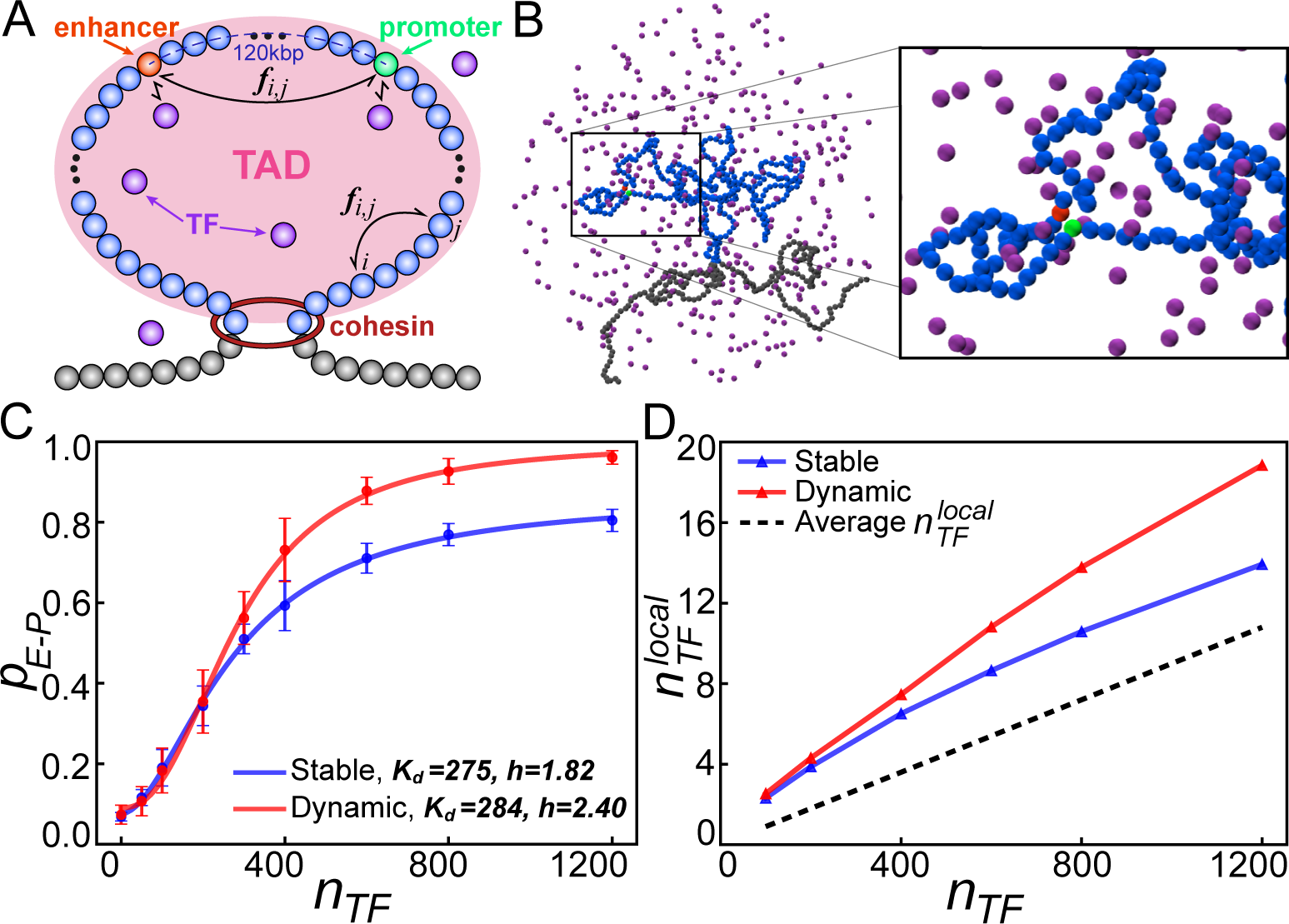
TF clustering in stable and dynamic chromatin models. (A) Schematic illustration of chromatin models in the presence of TFs (purple beads). This is similar to Figure 1A, with the addition of TFs in the system. TFs have attractive interactions only with the enhancer and promoter, modeling the specific interactions of TFs with DNA regulatory elements. (B) A simulation snapshot showing the formation of an E-P loop, with the chromatin chain surrounded by enriched TFs in 3D space. (C) E-P contact probability (*p_E−P_*) as a function of the number of TFs in the system (*n_TF_*) for stable and dynamic chromatin models. The data are fitted to the Hill function, with binding affinity *K_d_* and the Hill coefficient *h* displayed. Error bars represent the standard errors at the corresponding average values. (D) Number of TFs accumulating at the local E-P loci region 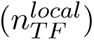 as a function of *n_TF_*. The dashed line indicates the average number of 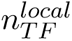 within the same volume of the local E-P loci region as if there is no chromatin in the system.

where *K_d_* is the effective binding affinity, *h* is the Hill coefficient, and *α*_0_ and *β* are other fitting parameters. We observed that *h* is greater than 1 for both models, indicating that the binding of TFs is cooperative in the formation of E-P contacts. Generally, a higher *h* indicates greater cooperativity among TFs in binding to the E-P loci.^55,56^ The larger *h* observed in the dynamic chromatin model suggests greater TF binding cooperativity and, consequently, more significant TF clustering at the E-P loci than in the stable model. Notably, since there are no attractive interactions between TFs, the observed binding cooperativity is entirely due to their interactions with the enhancer and promoter. Thus, the dynamic chromatin model can recruit more TFs to the E-P loci during the formation of E-P contacts compared to the stable model.

To quantitatively describe TF clustering, we introduced a metric to measure the degree of TF clustering: the number of TFs locally proximal to the E-P loci, denoted as 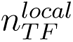 (details in Supporting Information). We found that when the number of TFs in the system (*n_TF_*) exceeds 400, the dynamic chromatin model presented a higher 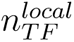, compared to the stable chromatin model, thus indicating a greater capacity for clustering TFs (Figure 2D). Meanwhile, the difference in 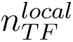 between the dynamic and stable models becomes more significant as *n_TF_* increases. This observation may be attributed to the broader conformational space and more significant fluctuations of chromatin led by the dynamic model, which can recruit more TFs to the proximity of E-P loci, compared to the stable model. Our findings are consistent with a recent study,^57^ suggesting that a flexible polymer can act as a chemical potential trap, facilitating polymer-assisted condensation. Therefore, we conclude that chromatin dynamics can induce the clustering of TFs, and this TF clustering, in turn, influences the chromatin structure by promoting persistent E-P contact, which can potentially contribute to the enhancement in transcriptional initiation.

To characterize the mechanism by which E-P communication is established, we quantified the free energy landscape for E-P contact formation. The free energy was calculated as: ℱ(*p_E−P_, d_E−P_*) = *−k_B_T* ln[𝒫(*p_E−P_, d_E−P_*)], where 𝒫(*p_E−P_, d_E−P_*) represents the probability of the system projected onto the reaction coordinates of E-P contact probability (*p_E−P_*) and E-P distance (*d_E−P_*) (details in Supporting Information). From the free energy landscape (Figure 3A), we observed three stable states during E-P contact formation: the P_on_ state, where E-P contact is successfully formed and is ready for subsequent gene expression initiation; the O_off_ state, where the enhancer and promoter are spatially distant; and the intermediate P_off_ state, where the E-P pair is spatially proximal but contact is not formed, which may not lead to transcriptional activation. A similar two-step E-P contact formation mechanism upon transcriptional activation has also been observed in experiments.^51,58^ Interestingly, we found that the intermediate P_off_ state disappears in the absence of TFs, and its stability increases as *n_TF_* increases (Figure S3). This result underscores the importance of TFs in stabilizing the P_off_ state.

**Figure 3:**
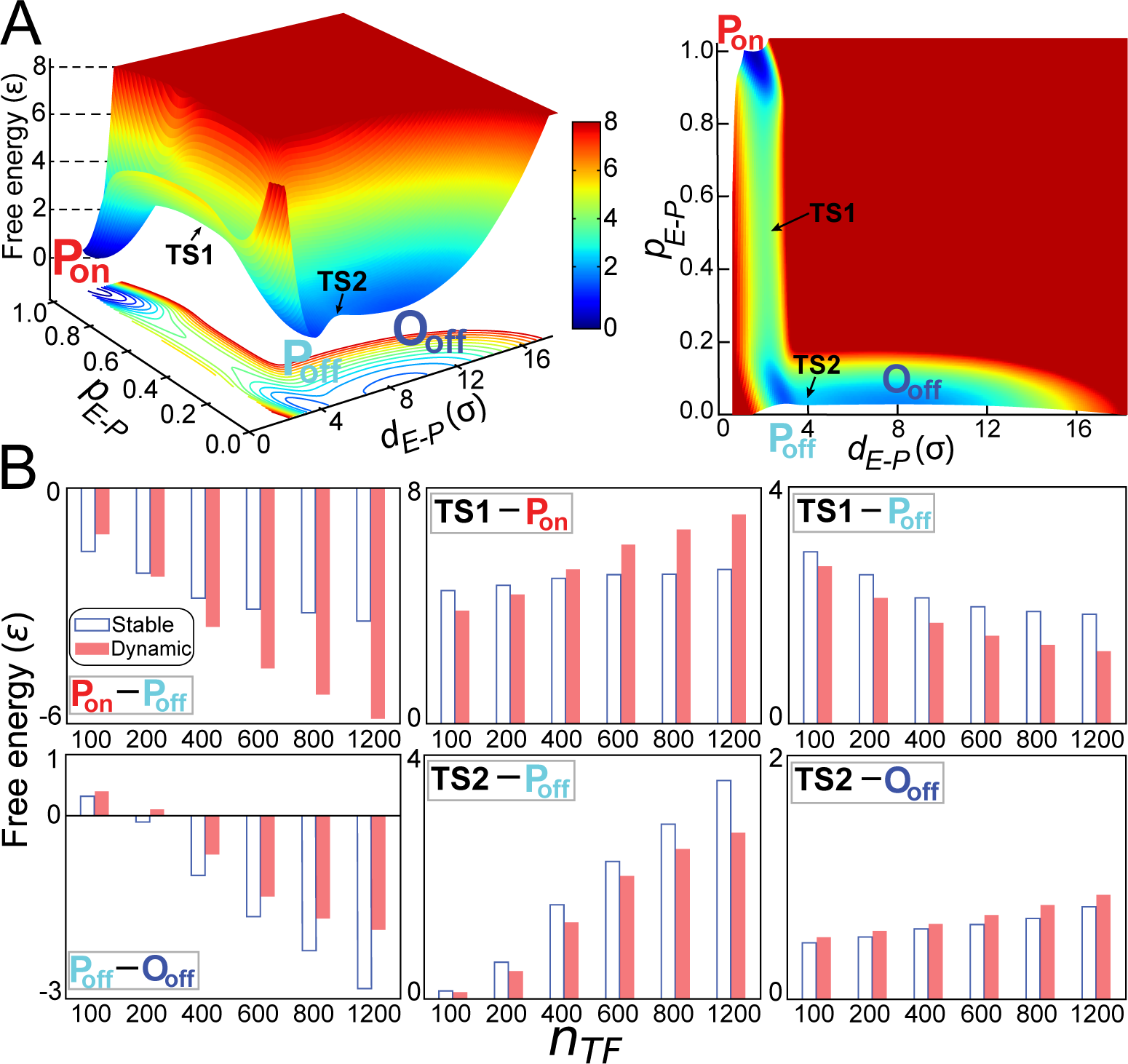
Free energy landscape and thermodynamic results of establishing E-P contact. (A) 2D free energy landscape projected onto *p_E−P_* and *d_E−P_*, when *n_TF_* = 200. Three stable states are observed: the P_on_ state, where the enhancer and promoter are spatially close and form a contact; the P_off_ state, where they are spatially close but do not form a contact; and the O_off_ state, where they are spatially far apart with no contact. The transition states between P_on_ and P_off_, and between P_off_ and O_off_, are indicated as *TS*1 and *TS*2, respectively. (B) Free energy differences between the stable states and the barrier heights during the state-transitions as a function of *n_TF_*.

To assess the effects of TFs on E-P contact formation, we calculated the free energy differences between the stable states (reflecting state stability) and between the stable states and transition states (TSs) (representing barrier heights) with increasing *n_TF_* for both the stable and dynamic chromatin models (Figure 3B, details in Supporting Information). We found that, for both models, the addition of TFs increases the stability of the P_on_ state relative to the P_off_ state. Additionally, the transition from the P_off_ state to the P_on_ state is accelerated with increasing *n_TF_*. Once the P_on_ state is formed, the E-P contact becomes more stable with increasing *n_TF_*, as evidenced by the increased barrier height between P_on_ to *TS*1. Notably, TFs have more significant impacts on the transitions between the P_on_ and P_off_ states by inducing more significant changes in stability and barriers for the dynamic model than the stable model.

On the other hand, TFs have little effect on the first transition step from the O_off_ state to the P_off_ state. However, the stability of the P_off_ state relative to the O_off_ state increases with increasing *n_TF_*, mainly due to the increase in the barrier height between *TS*2 and P_off_ when more TFs are added. Unlike the P_off_ to P_on_ state transition, the stabilization of the P_off_ state relative to the O_off_ state is less pronounced in the dynamic model compared to the stable model. Consequently, the dynamic chromatin model appears to disfavor the formation of the P_off_ state, compared to the stable model.

We further investigated the kinetics of E-P contact formation, which resembles the initiation of gene activation. We observed that the addition of TFs monotonically accelerates the formation of E-P contacts by reducing the mean first passage time (MFPT) for the transition from the O_off_ state to the P_on_ state (Figure 4A). Meanwhile, the dynamic chromatin model exhibited faster kinetics than the stable chromatin model at the same TF concentration. The effect of chain fluctuation on accelerating kinetics became more pronounced as *n_TF_*increased. To understand the reasons behind the differences in kinetics between these two models, we examined the TF-mediated clustering in E-P communication, represented by the TF crowding rate in the P_on_ and P_off_ states (Figure 4B, details in Supporting Information). The results revealed a significant enrichment of TFs between the enhancer and promoter in the P_off_ state compared to the P_on_ state. This crowding impedes the E and P from reaching a direct contact conformation due to volume exclusion between particles, further contributing to the stabilization of the intermediate P_off_ state. Interestingly, the dynamic chromatin model exhibited lower crowding rates in both the P_off_ and P_on_ states than the stable chromatin model. This is likely because the fluctuating chromatin chain can easily expel the TF cluster between the enhancer and promoter, thus facilitating the formation of E-P contacts, as evidenced from the analysis based on the barrier heights (Figure 3B).

**Figure 4:**
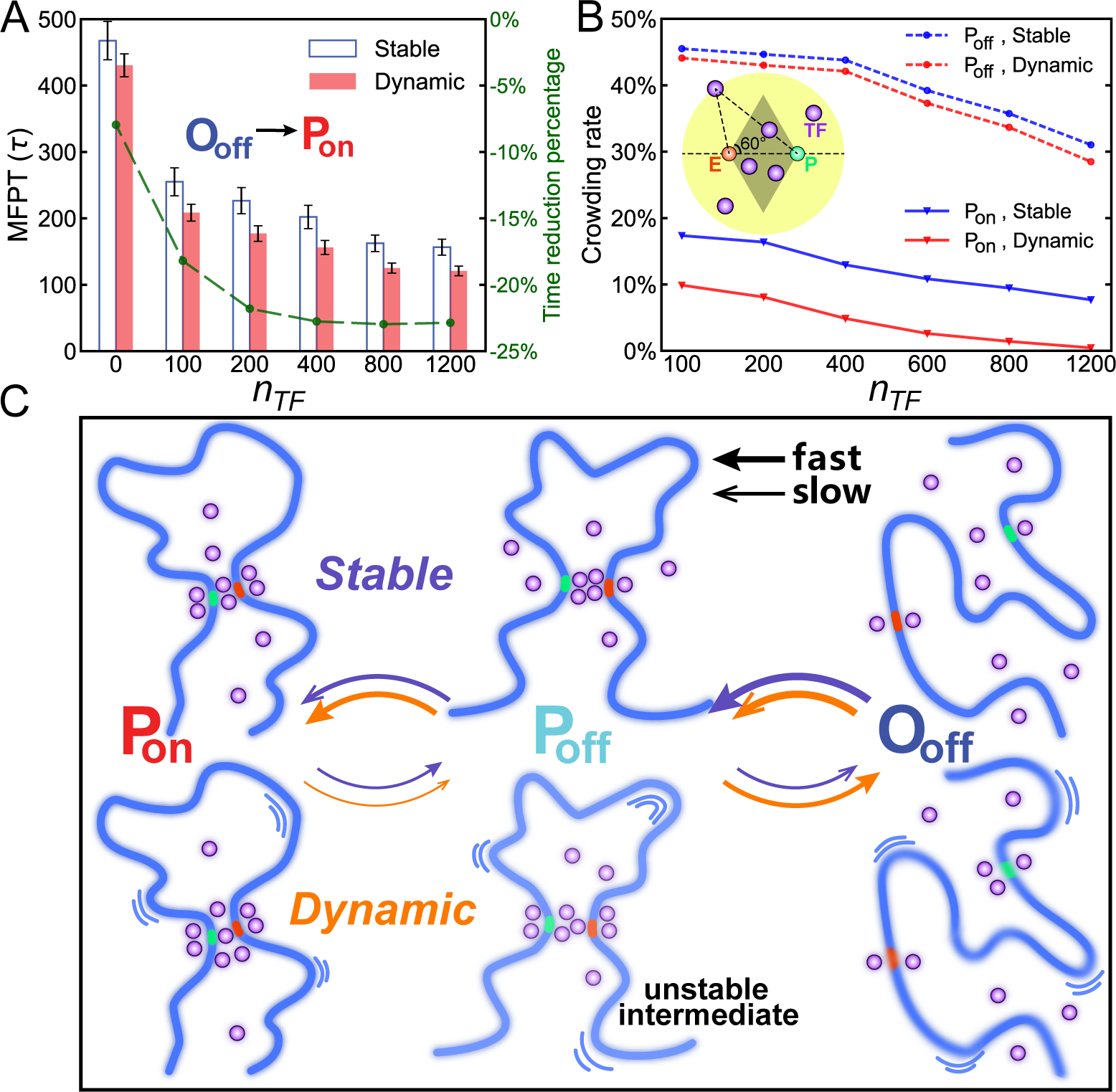
Kinetics of E-P contact formation modulated by TF clustering. (A) Mean first passage time (MFPT) for the transition from the O_off_ state to the P_on_ state as a function of *n_TF_* for the stable and dynamic chromatin models. Error bars represent the standard errors at the corresponding average values. The time reduction percentage was calculated as the ratio of the reduction in MFPT for the dynamic chromatin model compared to the stable chromatin model. (B) Quantitative description of TF clustering in terms of crowding rate as a function of *n_TF_*. The crowding rate is defined as the ratio of the number of TFs within the shaded dark diamond to the total number of TFs within the shaded yellow sphere, as indicated in the inset figure. The dark diamond and yellow sphere represent the regions mediating the E-P communication and regions proximal to the E-P loci, respectively (details in Supporting Information). (C) Schematic illustration summarizing the main findings of the study.

Together, we have elucidated the role of chromatin chain fluctuations in facilitating E-P contact formation through the modulation of TF clustering (Figure 4C). In the absence of TFs, forming E-P contacts in chromatin can be quite slow due to the lack of the intermediate P_off_ state. TF clustering, which occurs above certain TF concentrations, can promote the association of enhancer and promoter in spatial proximity, functionally serving as a transcriptional condensate.^30^ However, TF clustering also stabilizes the P_off_ state, which can hinder the formation of E-P contacts. For chromatin with significant structural fluctuations, the stability of the P_off_ state is relatively weak, allowing the system to easily escape from this state and undergo transition toward the P_on_ state, where the E-P contact is firmly established. Therefore, fluctuating chromatin facilitates E-P contact formation by efficiently inducing TF clustering and subsequently destabilizing the TF cluster, thereby promoting the formation of the transcriptionally active state.

Unraveling the dynamic properties of chromatin structures, which range from liquid-like to solid-like, is crucial for understanding the structure-function relationship in the genome.^34,59,60^ In this regard, we have demonstrated that similar population-averaged structural properties of chromatin can lead to significant differences in E-P communication, underlining the importance of chain fluctuations in regulating gene expression. Notably, chromatin dynamics are involved in various cellular functions, particularly cell development. During cell differentiation, chromatin motions within conserved TAD boundaries are progressively constrained.^61–66^ Dynamic chromatin structures in stem cells can promote high transcriptional activity to maintain pluripotency,^67^ echoing our theoretical findings, which suggest that chain fluctuations favor efficient E-P formation by dynamically modulating TF clustering without disturbing local chromatin architecture. Our study provides an advanced understanding of the role of chromatin dynamics in E-P communication and highlights the significance of chromatin structural fluctuations in gene regulation.

## Acknowledgement

X.C. acknowledges the support from the National Natural Science Foundation of China (Grant No. 32201020), the general program of Guangdong Basic and Applied Basic Research Foundation (Grant No. 2024A1515010862), the general program (Grant No. 2023A04J0083) and the Guangzhou-HKUST(GZ) joint funding program (Grant No 2023A03J0060) of the Guangzhou Municipal Science and Technology Project. X.C. was also partly supported by the Municipal Key Laboratory Construction program of the Guangzhou Municipal Science and Technology Project (Grant No. 2023A03J0003). C.L. was supported by the National Natural Science Foundation of China (Grant No. 12171102), and the National Key R&D Program of China (Grant No. 2019YFA0709502). The authors also acknowledge the Green e Materials Laboratory (GeM) and HPC+AI Intelligence Computing Center at the Hong Kong University of Science and Technology (Guangzhou) for providing computational support.

## Supporting Information Available

Supporting Information: Materials and methods, Figures S1-S9.

The necessary files for setting up the Gromacs simulations with PLUMED and the analysis programs/scripts are publicly available at https://github.com/icecolaTao/ChrModel. It includes the following components:

- Simulation Files: Gromacs input files for running the chromatin polymer model simulations.
- MEP Iteration Scripts: Python and Matlab scripts for performing MEP iterations.
- Analysis Tools: Python scripts necessary for analyzing simulation data.

This material is available free of charge via the Internet at http://pubs.acs.org/.

## Notes

### Competing Interest Statement

The authors have declared no competing interest.

