## Supplementary material for "Fluctuating chromatin facilitates enhancer-promoter communication by regulating transcriptional clustering dynamics": SI Text

### Supporting Information: Fluctuating chromatin domain facilitates enhancer-promoter communication by transcriptional clustering dynamics


Tao Zhu,† Chunhe Li,*,†,‡ and Xiakun Chu*,¶,§,‖

†Institute of Science and Technology for Brain-Inspired Intelligence, Fudan University, Shanghai 200433, China

‡Shanghai Center for Mathematical Sciences and School of Mathematical Sciences, Fudan University, Shanghai 200433, China

¶Advanced Materials Thrust, Function Hub, The Hong Kong University of Science and Technology (Guangzhou), Guangzhou, Guangdong 511400, China

§Guangzhou Municipal Key Laboratory of Materials Informatics, The Hong Kong University of Science and Technology (Guangzhou), Guangzhou, Guangdong 511400, China

‖Division of Life Science, The Hong Kong University of Science and Technology, Clear Water Bay, Hong Kong SAR 999077, China


### Materials and methods

#### Stable and dynamic chromatin models

The stable and dynamic chromatin models were generated by employing different non-bonded pairwise interaction potentials $f_{i,j}$. The stable model utilized the LJ potential $V_{\mathrm{LJ}}$, which, due to its attractive interaction nature acting on all pairs in the system, can significantly collapse the polymer chain, resulting in stable-like polymer structural ensembles. In the stable model, $V_{\mathrm{LJ}}$ is effective for any two beads $(i, j)$ that are separated by at least two beads, and is defined as:

$$V_{\mathrm{LJ}}\left(r_{i,j}\right) = 4\epsilon_{\mathrm{LJ}}\left[\left(\frac{\sigma_{LJ}}{r_{i,j}}\right)^{12} - \left(\frac{\sigma_{LJ}}{r_{i,j}}\right)^{6}\right] + \epsilon_{\mathrm{LJ}}.$$

We performed MD simulations using the stable chromatin model and calculated the Hi-C-like contact probability map $P_{i,j}^{Stable}$ for the structures obtained from the simulation trajectories. The contact probability $P_{i,j}$ was determined using the following expression:[1,2]

$$P_{i,j} = \frac{1}{2}\left[1 + \tanh\left[\mu\left(R_0 - r_{i,j}\right)\right]\right].$$

To generate a same contact map with highly heterogeneous and fluctuating structural properties, we applied the MEP method to construct the dynamic chromatin model.[1–3] The MEP approach allows for the creation of a model that produces a posterior distribution as close as possible to the prior distribution while satisfying the required observations (i.e., $P_{i,j}^{Stable}$). Consequently, the dynamic model, which uses the same bonded potential as the stable model but replaces the LJ potential with a data-driven non-bonded potential derived from MEP, can exaggerate the chromatin chain dynamics relative to the stable model.

In the dynamic model, two non-bonded potentials were used to replace the LJ potential from the stable model. The first term, $V_{sc}$, is a soft-core potential that allows partial overlap of the beads and is described by the following function:[1,2]

$$V_{\text{sc}}(r_{i,j}) = \begin{cases} 2\left(1+\tanh\left[0.5V_{\text{LJ}}\left(r_{i,j}\right)-1\right]\right), & r \leq r_0 \\ V_{\text{LJ}}\left(r_{i,j}\right), & r_0 < r_{i,j} \leq \sigma 2^{1/6} \\ 0, & r_{i,j} > \sigma 2^{1/6}, \end{cases}$$

where $r_0 = \sigma/[(1+\sqrt{2})/2]^{1/6}$.

The second term, $V_{\text{MEP}}$, is a data-driven potential restrained by the Hi-C-like contact map generated from the stable model and is expressed as:

$$\begin{aligned} V_{\text{MEP}}(r_{i,j}) &= \alpha_{i,j} P_{i,j} \\ &= \frac{1}{2}\alpha_{i,j}\left[1+\tanh\left[\mu\left(R_0 - r_{i,j}\right)\right]\right], \end{aligned}$$

where the prefactors $\alpha_{i,j}$ are determined through iterative simulations to gradually align the contact map of the dynamic model $P_{i,j}^{Dynamic}$ with that of the stable model $P_{i,j}^{Stable}$.

#### Simulation protocols and model parameters

We performed all MD simulations using Gromacs (version 4.5.7)[4] with the PLUMED plugin (version 2.5.0)[5] to implement spherical confinement. Reduced units were used throughout the simulations. Langevin dynamics was applied with a friction coefficient of $1.0\tau^{-1}$, where $\tau$ is the reduced time unit. The time step was set to $0.0005\tau$, and the simulation temperature, expressed in energy units, was set to $\epsilon$ unless otherwise specified. The diameter of the CG bead, $\sigma$, was set as the length unit.

For the bond potential, we set $r_0 = \sigma$ and $K_b = 3200\epsilon/\sigma^2$ to ensure that adjacent beads on the chain neither separate too far from each other nor overlap. The angle potential strength was set to $K_a = 2.0\epsilon$. For the non-bonded LJ potential, we set $\sigma_{LJ} = 1.5\sigma$ and used a uniform $\epsilon_{LJ} = 0.3\epsilon$ for interactions among all beads within the TAD, defining the stable model. The interaction between the enhancer and promoter was also described by the LJ potential with a stronger strength $\epsilon_{LJ} = 2.0\epsilon$, to model the binding specificity of E-P contacts. For the MEP potential, we used $\mu = 10.0\sigma$ and $R_0 = 2.5\sigma$, as suggested in the previous study.[1] The radius of the spherical confinement, $R_{\text{C}}$, was set to $32\sigma$ with a strength of $K_{\text{C}} = 100\epsilon/\sigma^2$. Non-bonded interactions were cut off at a distance of $7.5\sigma$.

The necessary files for setting up the Gromacs simulations with PLUMED and the analysis programs/scripts are publicly available at https://github.com/icecolaTao/ChrModel.

We further adjusted the strength of the LJ potential to $\epsilon_{LJ} = 0.15\epsilon$ in the stable model, keeping the other parameters unchanged, and then performed the same sets of simulations and analyses. Comparing these two stable models resulted in dense ($\epsilon_{LJ} = 0.3\epsilon$) and loose ($\epsilon_{LJ} = 0.15\epsilon$) chromatin structural ensembles, respectively (Figure S4). The loose chromatin model produced results similar to those of the dense chromatin model presented in the main text (Figures S4-S9), demonstrating the robustness of our findings, which are not heavily dependent on the specific parameter choices. However, we observed that the differences between the stable and dynamic chromatin models were less pronounced in the loose chromatin compared to the dense chromatin. This can be attributed to the fact that the loose chromatin model with LJ potential, may already exhibit a high degree of chain fluctuations, comparable to those induced by the MEP-derived model.

#### MEP simulations

To achieve adequate sampling during each iteration, we performed 120 independent simulations, each initialized from different chromatin structures. To further enhance sampling efficiency, we employed a two-stage simulation strategy, as described in previous studies.[2,6] In brief, the first stage involved simulated annealing, during which the temperature was linearly decreased from $2.0\epsilon$ to $\epsilon$ over a period of $500\tau$. The second stage consisted of a production simulation conducted at a constant temperature of $\epsilon$ for $2500\tau$. We collected all the structures from the last $2000\tau$ of each trajectory to calculate the contact probability $P_{i,j}$.

To evaluate the differences between the dynamic and stable models, we calculated the relative error $\Lambda$ between the contact probabilities derived from the dynamic model ($P_{i,j}^{\text{Dynamic}}$) and the stable model ($P_{i,j}^{\text{Stable}}$), which serves as the target for the MEP simulations:

$$\Lambda = \frac{\sum_{i,j} |P_{i,j}^{\text{Dynamic}} - P_{i,j}^{\text{Stable}}|}{\sum_{i,j} P_{i,j}^{\text{Stable}}}.$$

The optimization process was terminated when the relative error $\Lambda$ fell below 5% (Figure S1). Considering all non-bonded pairs in the system would require substantial computational resources and significantly slow down the MEP iteration process. To address this, we used a high-resolution index grid of 1:2:333 $\times$ 1:2:333 to filter contact pairs, as suggested in previous studies.[1,2,6] This approach is justified by the homogeneity of the system, except in the enhancer and promoter regions. For these specific regions, we employed a finer contact map, which includes all contact pairs for the beads within three-bead distances on either side of the enhancer or promoter, along with the enhancer and promoter beads themselves. After implementing these simplifications, the number of contact pairs in the system for MEP iterations was reduced to 27,584.

#### Simulations in the presence of TFs

For simulations involving TFs, we incorporated TFs into the chromatin system. These TFs interact with the enhancer and promoter through LJ potentials, while interactions among TFs, as well as between TFs and other chromatin beads, are modeled as non-specific volume-excluding effects. This approach is reminiscent of the strings-and-binders switch (SBS) model,[7] which has been successfully used to describe chromatin structural organization in the presence of TFs (binders) and has consistently yielded results that align with experimental observations.[8–10] The strength of the LJ potential between TFs and the enhancer and promoter was set to $\epsilon_{LJ} = 5.0\epsilon$. For the volume-excluding interactions among TFs and between TFs and other chromatin beads, the interaction is expressed as $4(\sigma/r_{i,j})^{12} + \epsilon$.

We performed simulations for both stable and dynamic chromatin in the presence of TFs, varying the $n_{TF}$ values: 0, 50, 100, 200, 300, 400, 600, 800, and 1200. For the stable chromatin simulations, we performed 40 independent simulations, each initialized from different system configurations. Each simulation lasted for $40,000\tau$, ensuring sufficient sampling, with data collected from the final $20,000\tau$ for analysis. Due to the slower simulation speed with the dynamic chromatin model, we shortened the length of each trajectory but increased the number of trajectories, ensuring that the total data collected was comparable to that of the stable model.

To examine how increasing the number of TFs affects E-P contact formation, we fitted the data points of $p_{E-P}$ and $n_{TF}$ using the Hill function, which has been widely applied in previous experimental studies:[11–13]

$$p_{E-P} = \alpha_0 + \frac{\beta n_{TF}^h}{K_d^h + n_{TF}^h}.$$

In this context, the Hill coefficient $h$ reflects the degree of binding cooperativity among TFs, with higher values indicating greater cooperativity or more TF clustering at the E-P loci. The dissociation constant $K_d$ represents the binding affinity of TFs to the E-P loci and is expressed in terms of the number of TFs.

To quantitatively characterize TF clustering, we defined $n_{TF}^{local}$ as the number of TFs concentrated at the E-P loci when the $d_{E-P}$ is sufficiently small. Specifically, when $d_{E-P}$ is less than $2.5\sigma$, we centered a sphere at the midpoint of the E-P axis with a radius of $7.5\sigma$ (as illustrated in Figure 4B of the main text, yellow region). The number of TFs within this sphere was then counted as $n_{TF}^{local}$. We calculated $n_{TF}^{local}$ for different total numbers of TFs ($n_{TF}$) present in the system.

#### Free energy landscapes and barrier heights of E-P interactions

We quantified the free energy landscape projected onto two coupled variables: the E-P distance ($d_{E-P}$) and the E-P contact probability ($p_{E-P}$) using the following expression:

$$\mathcal{F}(p_{E-P}, d_{E-P}) = -k_B T \ln[\mathcal{P}(p_{E-P}, d_{E-P})],$$

where $\mathcal{P}(p_{E-P}, d_{E-P})$ represents the probability at a given $d_{E-P}$ and $p_{E-P}$. The variable $d_{E-P}$ corresponds to the spatial distance between the enhancer and promoter, while $p_{E-P}$, derived from $d_{E-P}$ using a tanh function, primarily characterizes the interaction process when the enhancer and promoter are in close proximity. It has been established that $d_{E-P}$ effectively captures the unbound states with no contacts formed, whereas $p_{E-P}$ is more suitable for describing the process as contacts begin to establish.[14,15]

We then calculated the relative stability between the three stable states ($P_{on}$, $P_{off}$, $O_{off}$) and the barrier heights for transitions passing through the transition states TS1 and TS2 at various $n_{TF}$ values. Since direct calculations using the 2D landscapes can be cumbersome, we simplified the analysis by quantifying a 1D landscape using a single reaction coordinate, $\zeta_{E-P}$, which incorporates information from both $d_{E-P}$ and $p_{E-P}$ (Figure S3):

$$\zeta_{E-P} = d_{E-P} - 5 \times p_{E-P}.$$

From the 1D landscapes, we observed clear trends in barrier heights as $n_{TF}$ increases. Notably, when $n_{TF}$ is low (<100), the intermediate $P_{off}$ state is unstable. As $n_{TF}$ increases, the stability of the $P_{off}$ state also increases.

#### Calculations of MFPT and TF-crowding rate

To determine the mean first passage time (MFPT) for the transition from the $O_{off}$ state to the $P_{on}$ state, we performed additional simulations with various $n_{TF}$: 0, 100, 200, 400, 800, and 1200. For each value of $n_{TF}$, 100 independent trajectories were simulated, each initiated from a different configuration in the $O_{off}$ state. Each simulation was run for a duration of 4,000$\tau$. The first passage time (FPT) was defined as the time required for the system to reach the $P_{on}$ state for the first time. We then calculated the MFPT by averaging the FPT across all 100 trajectories.

To quantify the crowding of TFs between the enhancer and promoter, we introduced the TF-crowding rate. This metric is calculated as the number of TFs within the shaded region of Figure 4B divided by the number of TFs in the entire yellow spherical region (i.e., $n_{TF}^{local}$). The conical region (shaded area) is defined by an angle of 60° with respect to the E-P axis, encompassing the TFs that are within a distance $d_{E-P}$ from both the enhancer and promoter in the $P_{off}$ and $P_{on}$ states.

#### Physical units of model and simulation

In our study, the length unit $\sigma$ is defined as the diameter of each bead in the simulation, where each bead represents a 30-nm chromatin fiber, corresponding to a 3-kbp DNA segment.[16,17] Experimental study suggests that the persistence length of this chromatin fiber is between 170 and 220 nm,[18] leading to an estimate for $\sigma$ in the range of 170 to 220 nm.

A recent experimental study using live imaging to investigate chromatin dynamics in *Drosophila* examined the diffusive dynamics of an enhancer and promoter separated by a genomic distance of 149 kbp,[19] which is comparable to our simulation setting of 120 kbp. It was observed that the MSD of the promoter locus was approximately 0.6 $\mu m^2$ at a lag time of 100 seconds. By comparing our simulation results with this experimental data, we estimated $\tau$ to be in the range of 3.0 to 4.5 seconds.

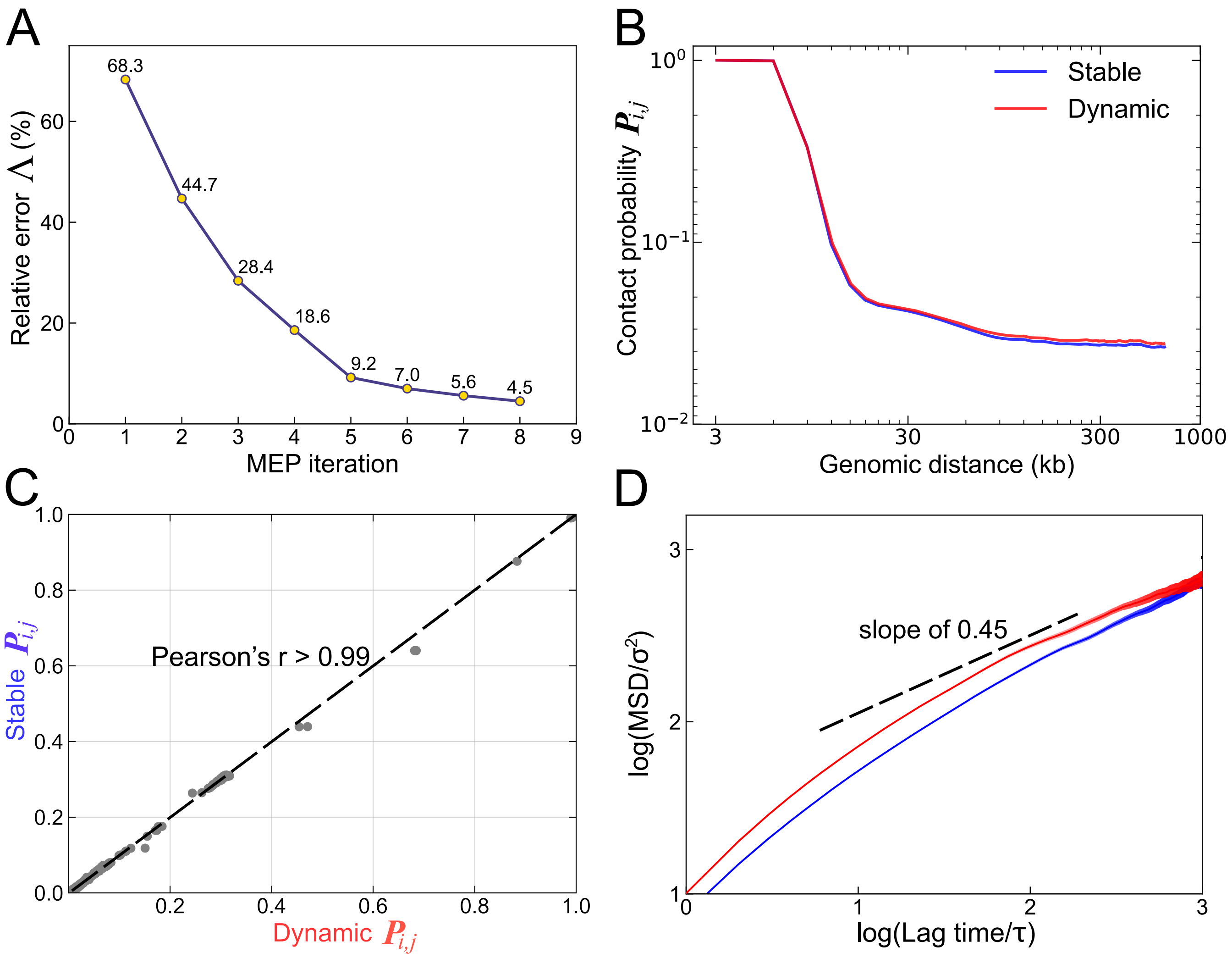


Figure S1: MEP simulations for generating the dynamic chromatin model.
(A) Relative error $\Lambda$ as a function of the MEP iterations. The MEP simulation was terminated after the $8^{\text{th}}$ iteration when $\Lambda$ dropped below 5%.
(B) Contact probability versus genomic distance for the stable and dynamic models.
(C) Correlation between the contact probabilities of all pairs of the 333 beads within the TAD for the stable and dynamic models.
(D) Logarithmic plot of the MSD of the promoter as a function of lag time for the stable and dynamic models.

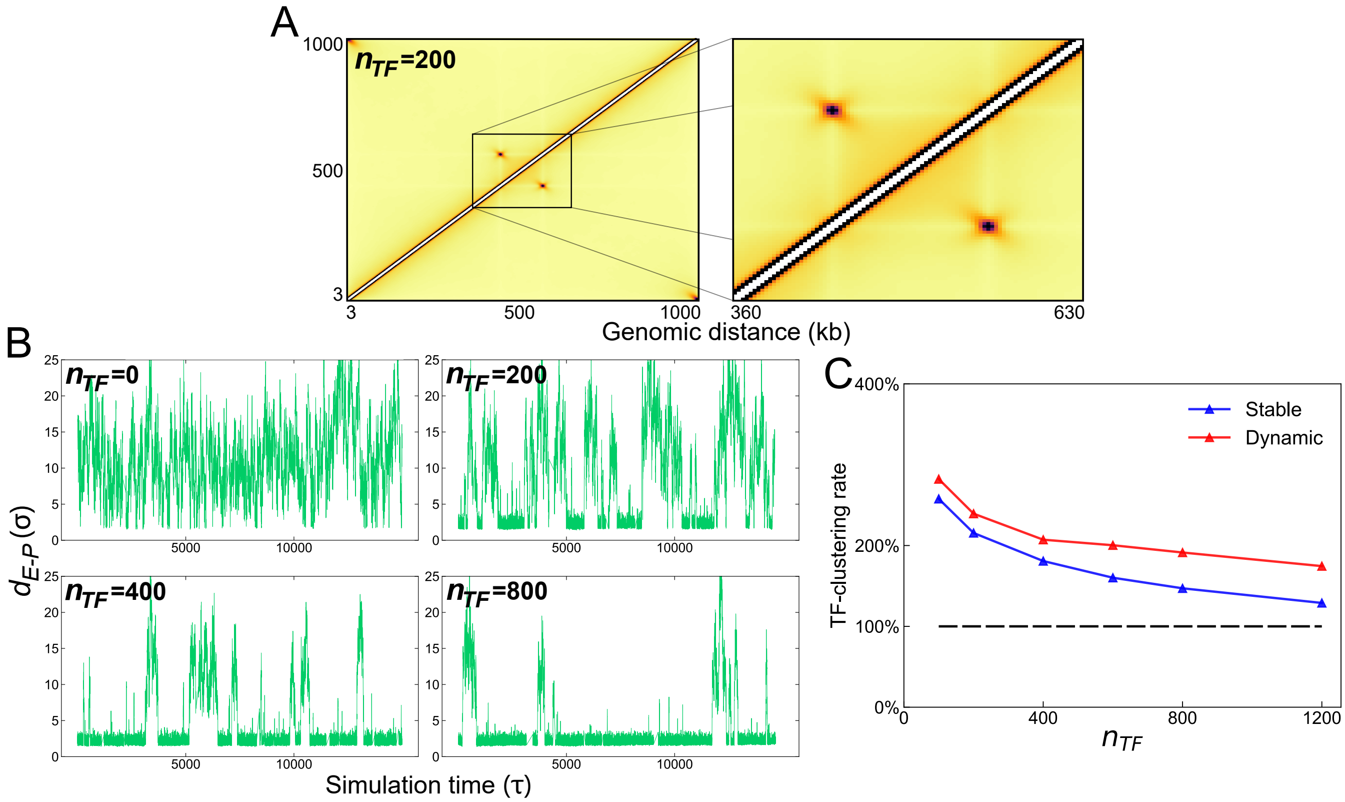


Figure S2: Contact maps, E-P distances and $n_{TF}^{local}$ in the presence of TFs.
(A) Hi-C-like contact maps for the stable chromatin model with $n_{TF} = 200$.
(B) Representative trajectories of $d_{E-P}$ for $n_{TF} = 0$, 200, 400, and 800.
(C) TF-clustering rate as a function of $n_{TF}$. The clustering rate is defined as $n_{TF}^{local}/n_{TF}(average)$. The denominator $n_{TF}(average)$ indicates the average number of $n_{TF}^{local}$ within the same volume of the local E-P loci region as if there is no chromatin in the system.

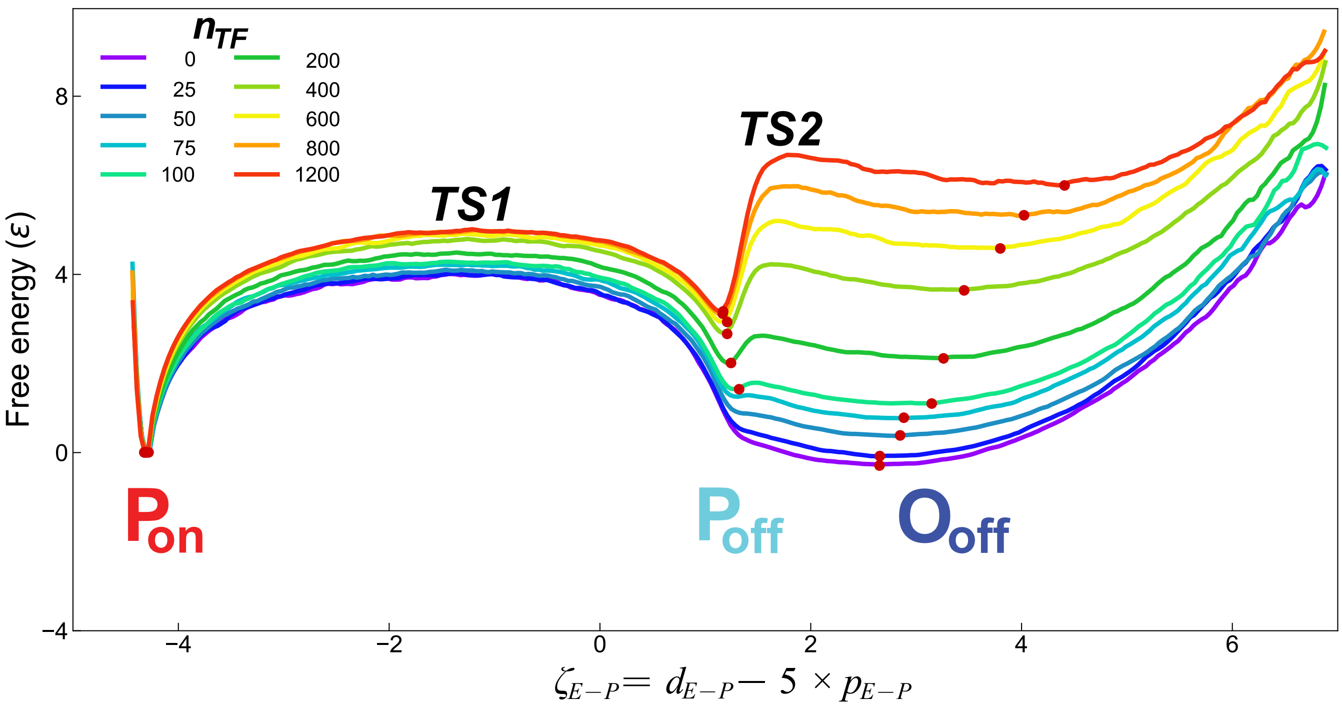


Figure S3: 1D free energy landscape projected onto the reaction coordinate $\zeta_{E-P}$.
The stable states ($P_{on}$, $P_{off}$, $O_{off}$) in the presence of certain numbers of TFs are clearly observed and indicated as red points, corresponding to the free energy minima.

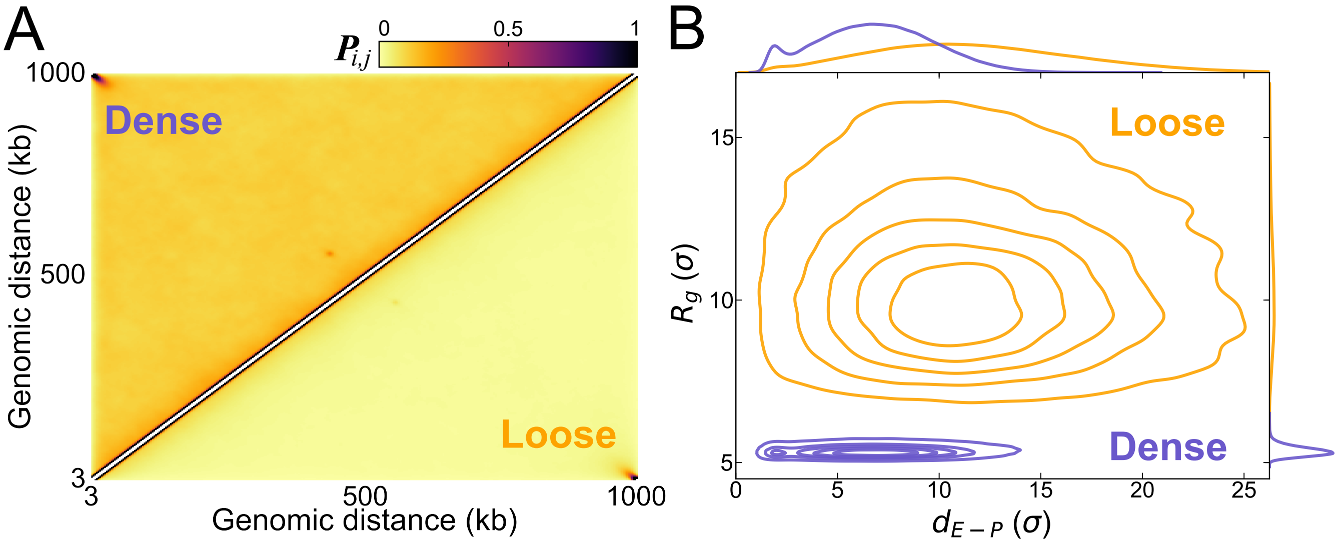


Figure S4: Structural properties of dense and loose chromatin models. Loose model is generated by applying $\epsilon_{LJ} = 0.15\epsilon$.
(A) Comparison between Hi-C-like contact maps for the dense and loose models.
(B) Distributions of the $R_g$ of the chromatin system and $d_{E-P}$ for the loose and dense stable models.

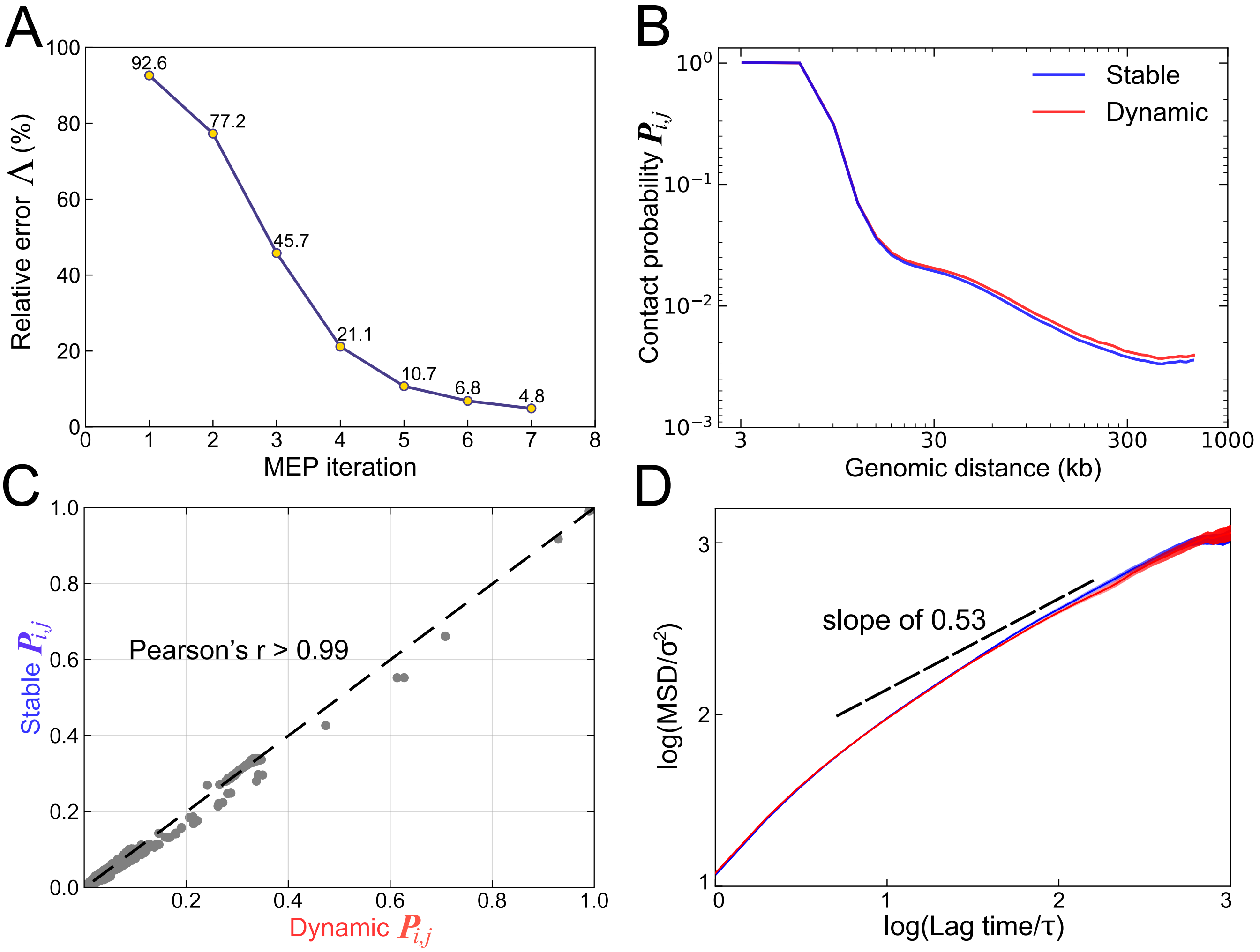


Figure S5: MEP simulation for generating the dynamic chromatin model from the loose stable chromatin model, where $\epsilon_{LJ} = 0.15\epsilon$.
(A-D) Similar with Figure S1A-D but for loose models.

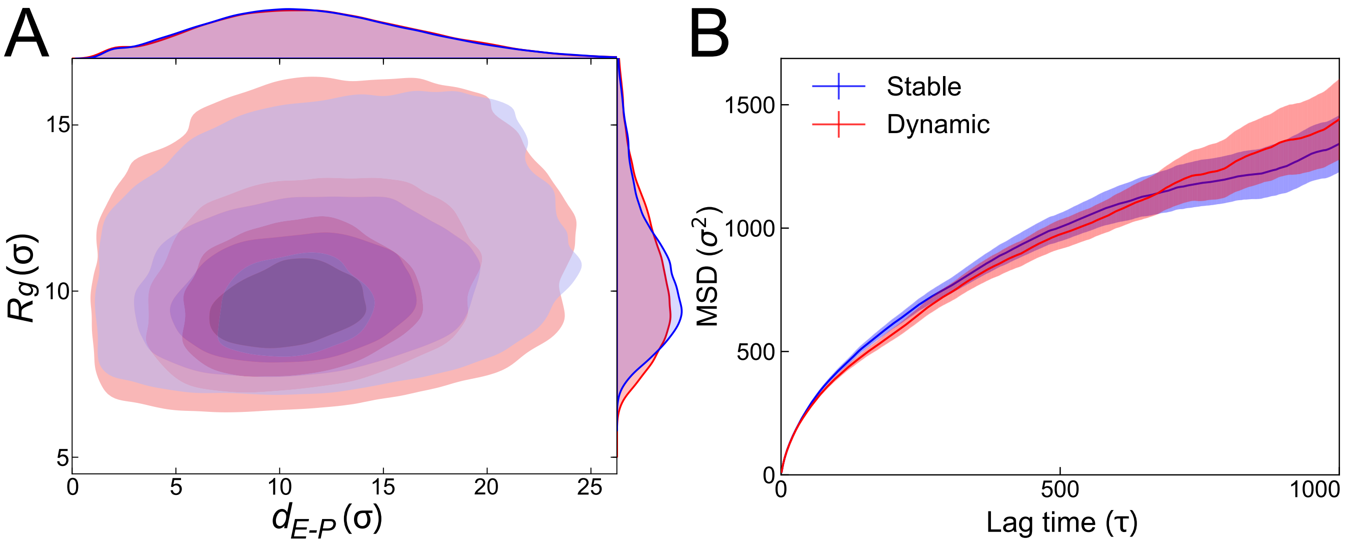


Figure S6: Structural and dynamic properties of loose stable and dynamic chromatin models, where $\epsilon_{LJ} = 0.15\epsilon$.
(A) and (B) Same with Figure 1E and 1F in main text, but for the loose model.

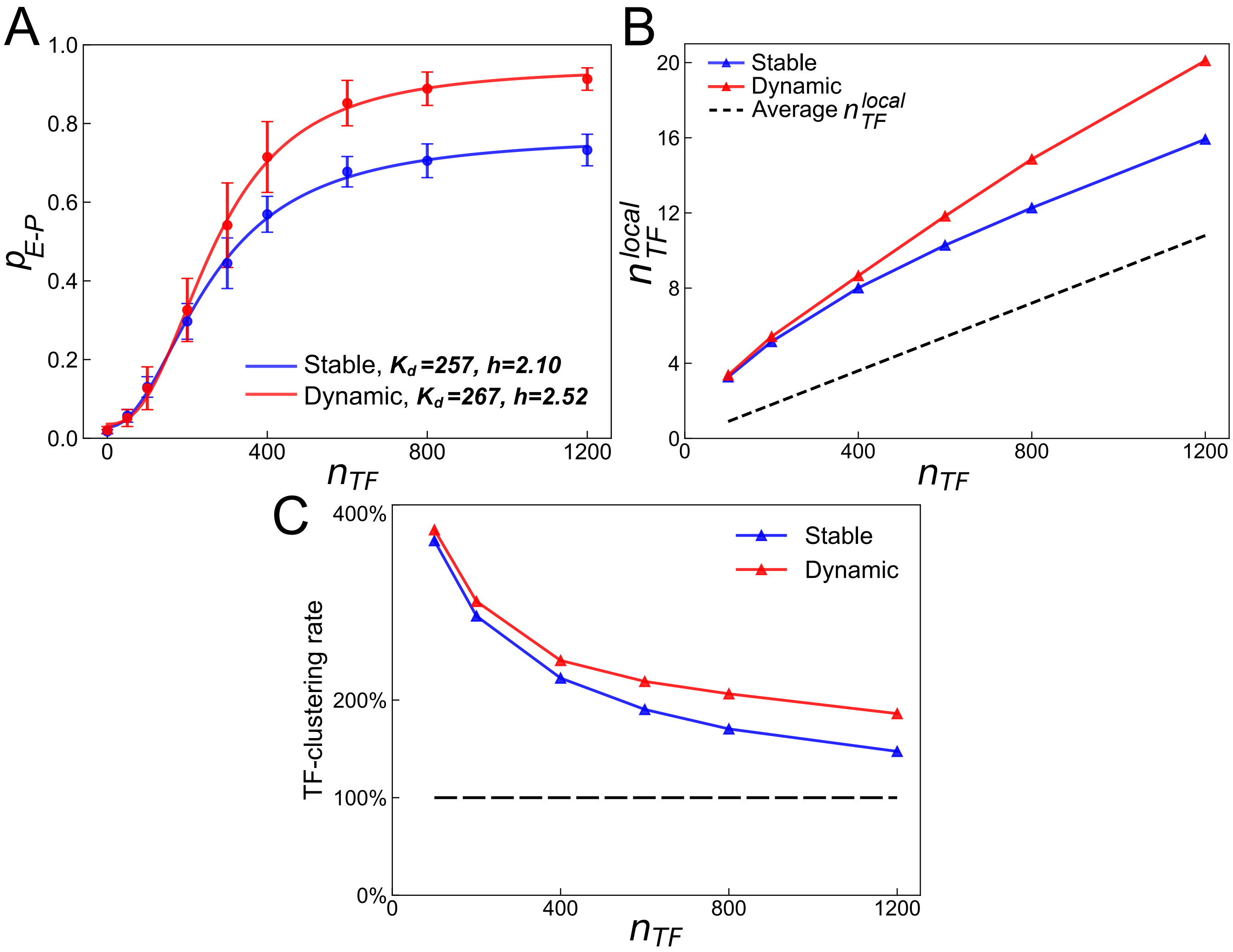


Figure S7: TF clustering in the loose stable and dynamic chromatin models.
(A) and (B) Same with Figure 2C and 2D in main text, but for the loose model.
(C) Same with Figure S2C, but for the loose model.

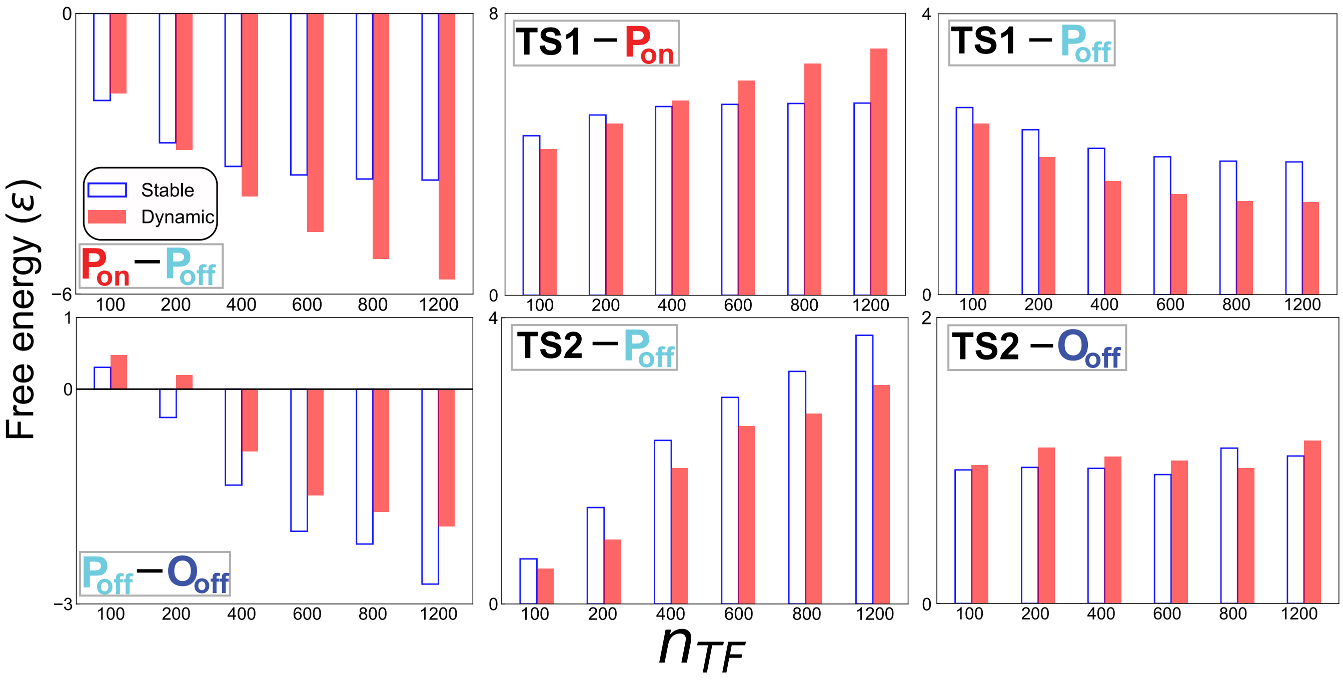


Figure S8: Thermodynamic results of establishing E-P contact for the loose stable and dynamic chromatin models.

The figure is the same with Figure 3B in the main text, but for the loose model.

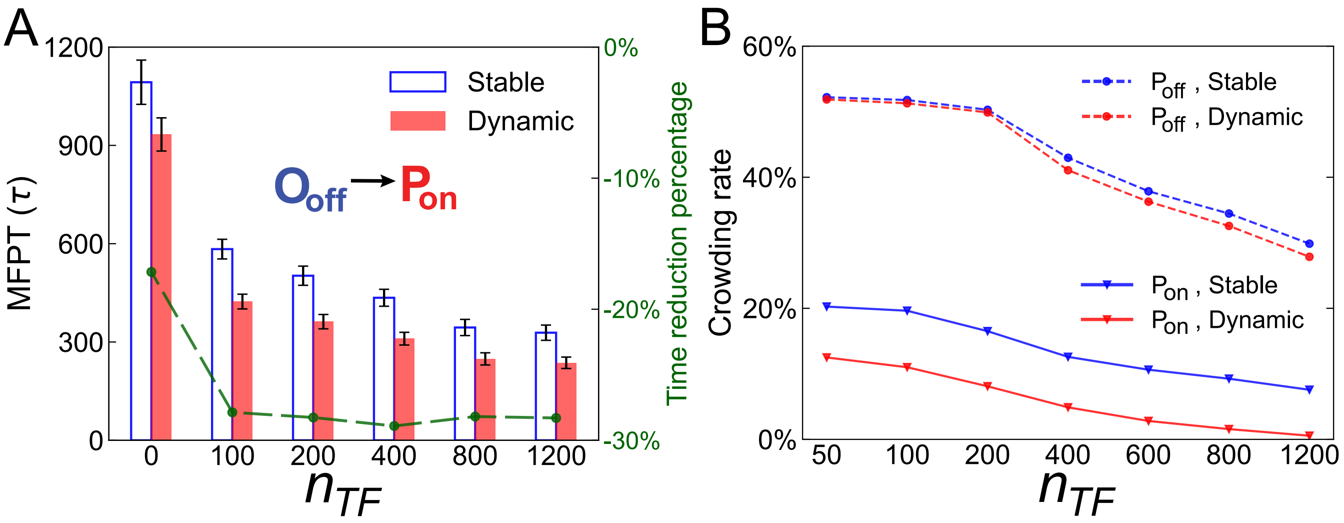


Figure S9: Kinetics of E-P contact formation modulated by TF clustering dynamics for the the loose stable and dynamic chromatin models.
(A, B) The same with Figure 4A and 4B in the main text, but for the loose model.

(16) Gerchman, S.; Ramakrishnan, V. Chromatin higher-order structure studied by neutron scattering and scanning transmission electron microscopy. *Proceedings of*

*the National Academy of Sciences* **1987**, *84*, 7802–7806.

(17) Brackley, C. A.; Johnson, J.; Kelly, S.; Cook, P. R.; Marenduzzo, D. Simulated binding of transcription factors to active and inactive regions folds human chromosomes into loops, rosettes and topological domains. *Nucleic acids research* **2016**, *44*, 3503–3512.

(18) Bystricky, K.; Heun, P.; Gehlen, L.; Langowski, J.; Gasser, S. M. Long-range compaction and flexibility of interphase chromatin in budding yeast analyzed by high-resolution imaging techniques. *Proceedings of the National Academy of Sciences* **2004**, *101*, 16495–16500.

(19) Brückner, D. B.; Chen, H.; Barinov, L.; Zoller, B.; Gregor, T. Stochastic motion and transcriptional dynamics of pairs of distal DNA loci on a compacted chromosome. *Science* **2023**, *380*, 1357–1362.
